# Correction of z-motion artefacts to allow population imaging of synaptic activity in awake behaving mice

**DOI:** 10.1101/767640

**Authors:** Thomas Ryan, Antonio Hinojosa, Rozan Vroman, Christoforos Papasavvas, Leon Lagnado

## Abstract

Functional imaging of head-fixed, awake, behaving mice using two-photon imaging of fluorescent activity reporters has become a powerful tool in the studying the function of the brain. Motion artefacts are an inevitable problem during such experiments and are routinely corrected for in x and y dimensions. However, axial (z) shifts of several microns can also occur, leading to intensity fluctuations in structures such as synapses that are small compared to the axial point-spread function of the microscope. Here we present a simple strategy to correct z-motion artefacts arising over the course of a time-series experiment in a single optical plane. Displacement in z was calculated using dye-filled blood vessels as an anatomical marker, providing high contrast images and accuracy to within ∼0.1 µm. The axial profiles of ROIs corresponding to synapses were described using a Moffat function and this “ROI-spread function” used to correct activity traces on an ROI-by-ROI basis. We demonstrate the accuracy and utility of the procedures in simulation experiments using fluorescent beads and then apply them to correcting measurements of synaptic activity in populations of vasoactive-intestinal peptide (VIP) interneurons expressing the synaptic reporter SyGCaMP6f. Correction of z-motion artefacts had a substantial impact on the apparent correlation between synaptic activity and running speed, demonstrating the importance of correcting for these artefacts for the interpretation of *in vivo* imaging experiments in awake mice.

**Summary of Key Points:**

- Motion artefacts associated with motor behaviour are an inevitable problem of multiphoton imaging in awake behaving animals, particularly imaging synapses.
- Correction of axial motion usually requires volumetric imaging resulting in slower rates of acquisition.
- We describe a method that is easy to implement to correct z-motion artefacts that allows population imaging of synaptic activity while scanning a single plane in a standard multiphoton microscope.
- The method uses a reference volume acquired in two colour channels – an activity reporter and an anatomical marker of blood vessels. The procedure estimates the z-displacement in every frame and applies an intensity correction in which the z point-spread function for each synapse is modelled as a Moffat function.
- We demonstrate that the method allows synaptic calcium activity signals to be collected from populations of synaptic boutons in mouse primary visual cortex during locomotion.

## Introduction

Many of the functions of the nervous system can be broken down into tasks carried out by sets of neurons connected through synapses into circuits. Investigating the operation of these circuits has been greatly advanced by using multiphoton microscopy to image fluorescent reporter proteins that signal events such as changes in cytoplasmic calcium, reflecting spiking activity (Tian *et al*., 2012), or the release of glutamate at excitatory synapses (Marvin *et al*., 2013). At present, the large majority of *in vivo* imaging studies quantify activity as calcium signals in the soma of neurons but we also need to image activity across populations of synapses if the operations of neural circuits are to be unravelled (Lu *et al*., 2017; Kazemipour *et al*., 2019; Meng *et al*., 2019). These connections are key components of all the computations within neural circuits and sites at which neuromodulators act to reconfigure signal flow according to changes in the internal state of the animal. The development of genetically encoded activity reporters that are targeted to synapses has been a crucial step towards understanding the biological machinery underlying neural computations. For instance, SyGCaMPs are localized to synaptic vesicles and report the calcium transients in synaptic boutons that activate neurotransmitter release (Miesenbock *et al*., 1998; Dreosti *et al*., 2009; Broussard *et al*., 2018; Schröder *et al*., 2019) while iGluSnFR expressed on the cell surface reports extracellular glutamate and can be used to detect individual vesicles (Marvin *et al*., 2013, 2018; James *et al*., 2019).

A fundamental issue in analysing *in vivo* imaging experiments in awake mice is the problem of movement of the brain while the animal is under the microscope. Such movements can be caused by breathing, locomotor behaviour or actions such as licking to obtain a reward. To correct motion artefacts in the x-y dimensions images can be co-registered to realign compartments before quantifying the signal in each, the most common algorithms making a series of rigid or non-rigid shifts of each image (Greenberg & Kerr, 2009; Pachitariu *et al*., 2016; Pnevmatikakis *et al*., 2016). Motions of the brain along the z-axis of the microscope can also occur, causing regions of interest (ROIs) to change their intensity as they go in or out of focus (Dombeck *et al*., 2007). In practise, the majority of two-photon imaging studies in awake animals do not correct for z-motion because calcium signals are usually acquired from the soma of neurons, which are ∼10 µm in diameter and therefore change intensity relatively little when displaced a few microns. But most synaptic boutons in the mouse cortex are <1 µm in diameter and appear considerably brighter or dimmer during z-motion of a few microns, confounding attempts to monitor activity in the awake animal, especially if that activity is itself correlated with locomotion or other behaviour of the mouse. Artefacts arising from z-motion are less straightforward to correct than lateral motion (Harris *et al*., 2016), requiring knowledge of both the displacement in z and the function relating this displacement to a change in intensity.

How can z-motion artefacts be dealt with to image synaptic activity with accuracy? One strategy is to reduce the artefact by elongating the point-spread function (PSF) of the microscope through the creation of Bessel beams (Garces-Chavez *et al*., 2002) or, more simply, by reducing the numerical aperture (NA) of the illumination beam. Both approaches, however, come at the cost of a substantial decrease in signal-to-noise ratio (SNR) due to increased background signal and decreased efficiency of two-photon excitation. A second general approach is to use volumetric imaging, in which multiple planes are scanned cyclically through a volume that contains all likely z-positions of the compartments of interest allowing realignment in three dimensions. But volumetric imaging introduces a trade-off: the larger the number of planes the more accurate the correction but at the cost of reduced time resolution. The continuing development of activity reporters with faster kinetics makes it increasingly important to image at the highest frequencies possible (James *et al*., 2019; Kazemipour *et al*., 2019). A third strategy might be to correct for z-motion by deconvolution of images in the time-series using the PSF of the microscope measured in 3D. This does not work well in practise because of the optical aberrations introduced by the cranial window and the brain tissue, which vary as a function of x-y-z position in the brain (Ji *et al*., 2012). Correcting local aberrations has required specialized adaptations of a multiphoton microscope such as the introduction of spatial light modulators into the light path to implement adaptive optics (Ji *et al*., 2012; Ji, 2017; Meng *et al*., 2019).

Here we describe a relatively simple approach to the correction of z-motion artefacts when imaging synaptic activity in awake behaving mice. The method can be used when imaging in a single plane on any standard commercial multiphoton microscope without the need for further specialised equipment. Estimates of z-motion are made from simultaneously obtained images of an inactive anatomical marker. Rather than using a second fluorescent protein, we use a dye that fills blood vessels, providing brighter images of greater spatial contrast and, therefore, improved accuracy. Knowledge of the absolute motion is then used to correct intensity measurements within each ROI by taking into account the effective PSF measured for each synapse individually. An important aspect of this correction is to reduce noise by modelling the PSF as a Moffat function rather than the more usual Gaussian, an approach similar to that used by astronomers imaging stars through atmospheric turbulence (Moffat, 1969; Racine, 1996; Starck *et al*., 2007). Using population synaptic recordings in mouse primary visual cortex (V1), we demonstrate how this approach disambiguates physiological activity from z-motion artefacts arising from locomotor behaviour.

## Methods

### Animals and surgical procedures

All experimental procedures were conducted according to the UK Animals Scientific Procedures Act (1986). Experiments were performed at University of Sussex under personal and project licenses granted by the Home Office following review by the Animal Welfare and Ethics Review Board (AWERB).

Experiments were performed on three *VIP-Cre* mice (Taniguchi *et al*., 2011), two females and one male, maintained on a C57BL/6J background. At post-natal day 60-90, surgery was performed to implant a cranial window over the primary visual cortex, similarly to described elsewhere (Holtmaat *et al*., 2009). In brief, dexamethasone (2 mg/ml) was administered by subcutaneous injection approximately 12 hr and 3 hr prior to surgery to reduce inflammation and brain swelling. Anaesthesia was induced and maintained by isoflurane inhalation (5% for induction, 0.5-2% maintenance). Analgesic (Meloxicam, 0.2 mg/kg) was administered subcutaneously after induction. During surgery, a custom-built titanium headplate was attached to the skull using dental cement (Unifast Trad; GC America, Chicago, IL, U.S.A.). A circular craniotomy 3 mm in diameter (centred 3 mm lateral and 0.5 mm anterior of Lambda) was drilled by hand over left monocular V1 and the edges of the craniotomy were bevelled to allow fixation of a cranial window. Prior to implantation of the window, a viral preparation was injected into V1 to drive expression of SyGCamMP6f in a Cre-recombinase dependent manner (AAV9 phSyn1(S)-FLEX-Sy-GCaMP6f-WPRE, ∼8 × 10^12^ viral genome copies/ml) using a stereotactic frame and injection apparatus (UMP3 UltraMicroPump, Nanofil syringe; WPI, Sarasota, FL, USA). In one female mouse, this was co-injected with a second virus driving expression of tdTomato in the same cells (AAV9 CAG Flex tdTomato WPRE, ∼ 2 × 10^12^ viral genome copies/ml). Typically, we injected 250 nl at a rate of 25 nl/min in three locations at a depth of 300 µm targeting L2/3. Windows consisted of three circular 3 mm coverslips with a 5 mm coverslip on top (No. 1 thickness; Menzel-Glaser, Braunschweig, Germany), all fixed together using optical glue (NOI 61; Norland Products, Cranbury, NJ, USA). Once implanted and sealed with dental cement, buprenorphine (0.02 mg/kg) was administered intramuscularly and anaesthesia terminated. Mice were administered meloxicam (0.3 mg/kg body weight per day) in wet mash food for 4 days following surgery as post-operative analgesic.

### Two-photon imaging

Imaging in V1 was carried out in head-fixed mice using a Scientifica two-photon microscope (Scientifica, Uckfield, East Sussex, U.K.) equipped with a mode-locked Chameleon titanium–sapphire laser tuned to 915 nm (Coherent Inc., Santa Clara, CA, U.S.A.) with an Olympus XLUMPLFLN 20× water immersion objective (NA 1, Olympus, Tokyo, Japan). Scanning and image acquisition were controlled under ScanImage v.3.8 software (Pologruto *et al*., 2003). The microscope was equipped with a piezo-controlled objective mount (PiFoc; Physik Instrumente, Karlsruhe, Germany) and a LabJack T7 module (Labjack, Lakewood, CO, U.S.A.) for Input/Output utility. Two channel recordings of SyGCaMP6f, tdTomato or dextran-TexasRED in L2/3 (180-300 µm depth) were taken simultaneously. Images were typically captured at a zoom factor of 7, an x,y, resolution of 512 × 200 pixels and a frame rate of 10.8 Hz. Visual stimuli were delivered through two computer monitors (Dell, Bretford, Middlesex, UK) and generated using the PsychoPy toolbox (Peirce, 2007). Animals were free to run on a cylindrical polystyrene treadmill (22 cm circumference, 7cm width). Movement of the treadmill was signalled by a rotary encoder (2,400 pulses per rotation, Kübler, Villingen-Schwenningen, Germany) and streamed through the LabJack using custom written software in MatLab (MathWorks, Natick, MA, U.S.A.), into the same Data Acquisition (DAQ) Board collecting images (National Instruments, Berkshire, Newbury, UK). The z-position of the objective was detected through a position sensor within the objective mount and recorded through the same DAQ recording images through ScanImage software. This strategy conveniently synchronised the acquisition of images through the photodetectors with signals from the rotary encoder and objective mount: data streams from these different sources were saved into a single multi-channel image file with TIFF format. Recordings from each mouse were acquired over multiple days over a 3 month period. Each field of view typically contained 20-50 active synapses.

### Simulations with fluorescent beads

Green fluorescent beads (FluoSpheres 1 µm diameter blue-green fluorescent; Thermo Fisher, London, UK) were sonicated for 2 mins, before preparing a 1:3000 dilution in 2% low-melting point agarose, a drop of which was placed on a deep welled slide which was then covered by three coverslips, as used for the cranial window of mice. Beads were imaged in the same way as neurons in V1 and controlled displacements of the focal plane imposed through the piezo-mount of the objective. These displacements reproduced those recorded during mouse locomotion.

### Labelling of blood vessels

To obtain images of the structure of V1 independent of activity, a water-soluble dextran conjugated fluorescent dye (Dextran, Texas Red, 3000 MW; Thermo Fisher, London, UK) was injected subcutaneously prior to an imaging experiment. Generally, a good fluorescence signal in brain capillaries was observed after ∼10 mins and remained for ∼2.5 hr, before fading over ∼30 mins. Imaging sessions always began more than 20 mins after injection and concluded within 1.5 hr, ensuring consistent blood vessel intensity for the whole session. Comparisons with the fluorescent protein marker of neurons tdTomato demonstrated that visualization of blood vessels in this way allowed significantly more accurate estimates of z-motion (Fig. 4).

**Figure 1.**
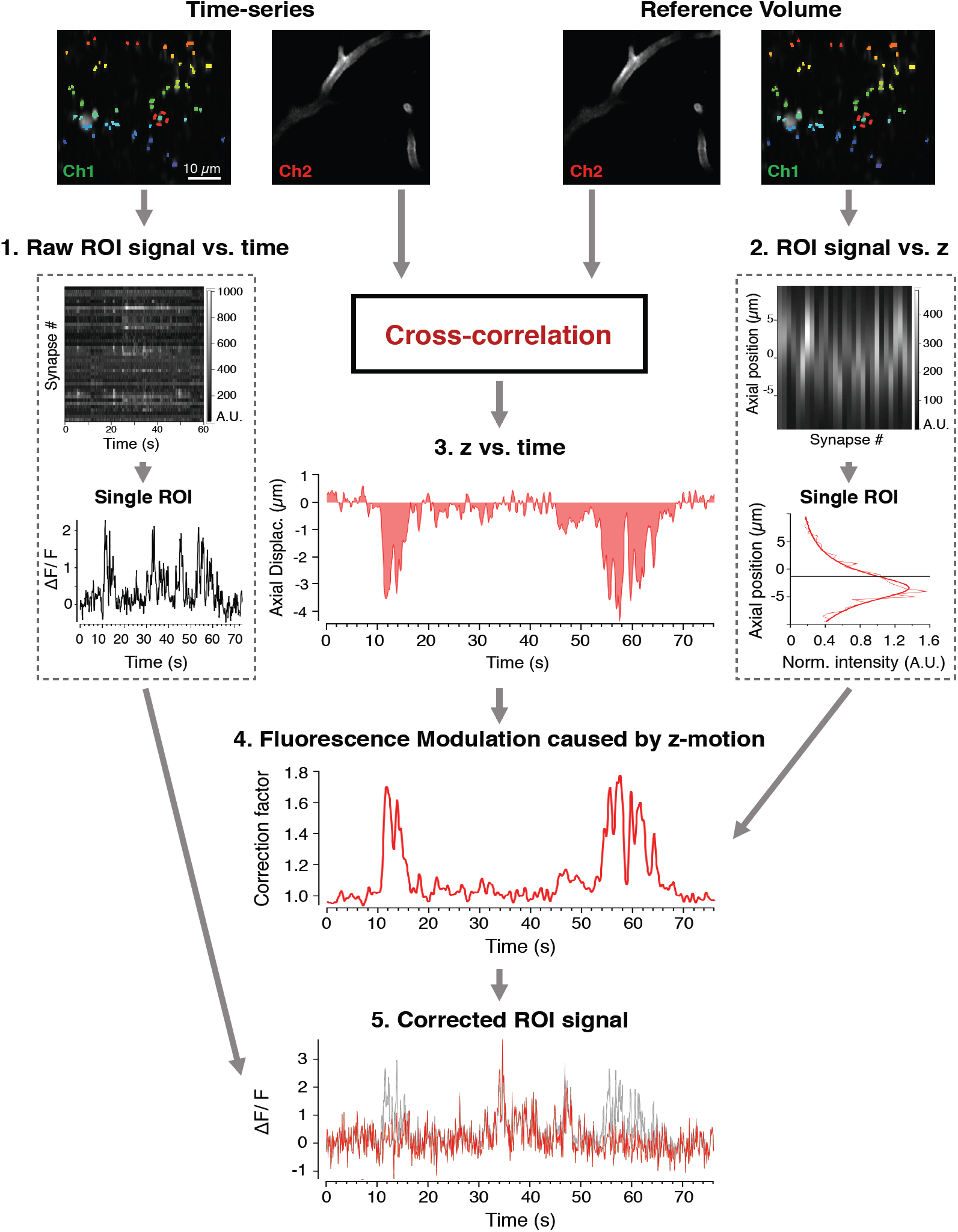
Workflow for correction of z-motion artefacts when imaging synaptic activity *in vivo*. A two-channel time-series (top left) and reference volume (top right) is acquired; SyGCaMP6f, green channel (Ch1); blood vessel marker, red channel (Ch2). An ROI mask is generated and applied to Chl of the time-series (coloured pixels). The raw signal over time is collected for all ROIs. This relation is shown in box 1, where the upper panel shows each ROI represented as a row and the bottom panel shows an example of an individual trace. The same ROIs are identified in the reference volume, providing a function relating ROI intensity with z-position (box 2). The continuous line describing these measurements is a Moffat Function (equation 1) with α =3.83 and β=1.33. The axial position as a function of time (relation 3) is estimated by cross-correlation between each frame in the time-series and a reference volume in Ch2. Relations 2 and 3 are then used to calculate a correction vector (relation 4) that estimates the relative change in fluorescence caused by z-motion through the time-series. The signal from each ROI can then be divided by its own correction vector to obtain an estimate of signal independent of z-position (relation 5, original signal in grey, corrected in red).

**Figure 2.**
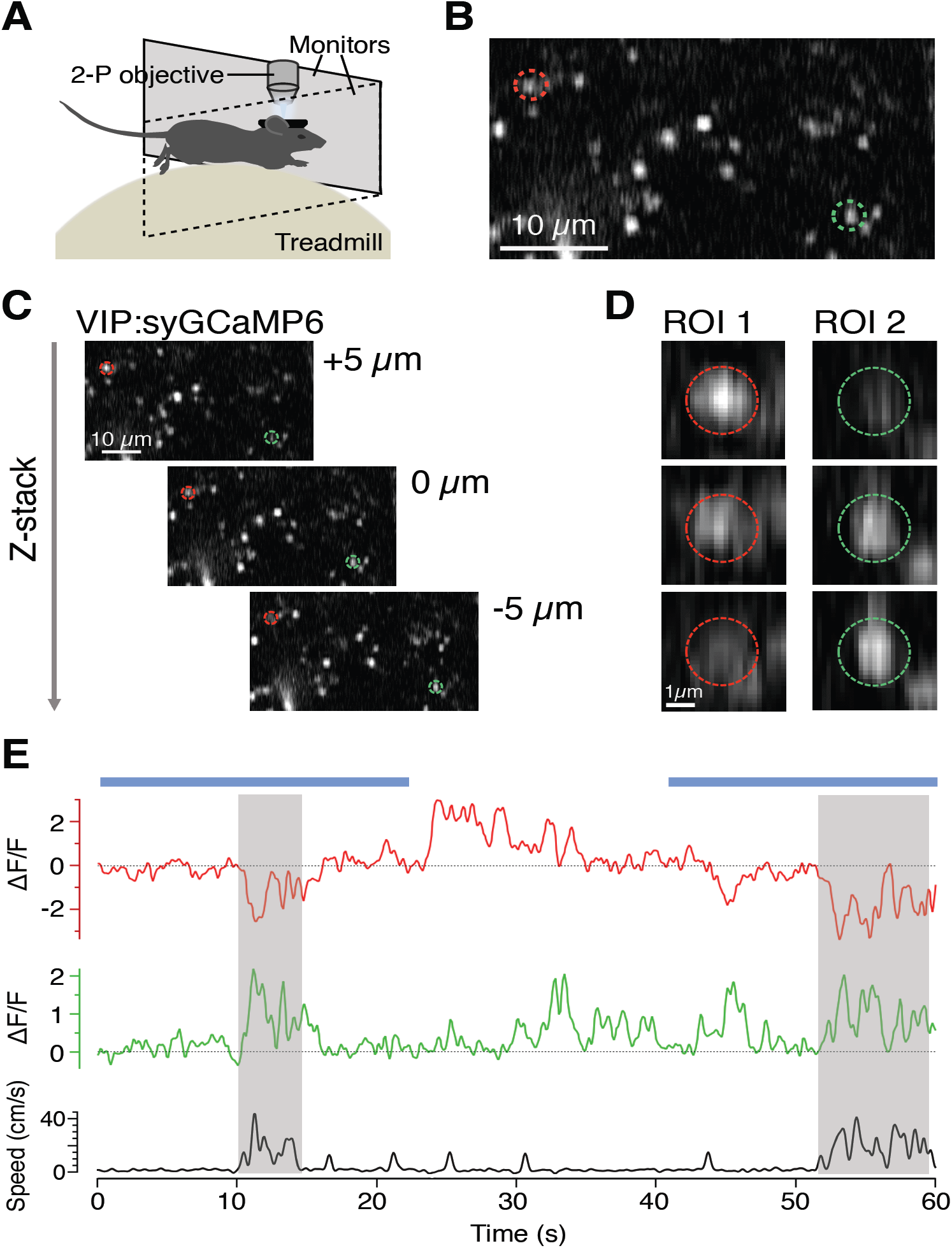
Imaging synaptic activity in awake behaving mice. **A**. Basic experimental set up for *in vivo* imaging. Awake animals are head-fixed under two-photon (2-P) excitation while the animal is allowed to run freely on a polystyrene treadmill connected to a rotary encoder to record running speed. Visual stimuli are delivered through two computer monitors placed in front of the animal. **B**. Example two-photon image of a small field of synapses labelled with SyGCaMP6f. **C**. Two-photon images of the same field of synapses at imaging depths separated by 5 µm intervals. **D**. The synapses circled in B and C exhibit significant changes in intensity when the focal plane changes over a few microns. **E**. Uncorrected fluorescence time-series compared to locomotor behaviour for the same synapses. Blue bars show the timing of a stimulus (fullfield drifting grating).

**Figure 3.**
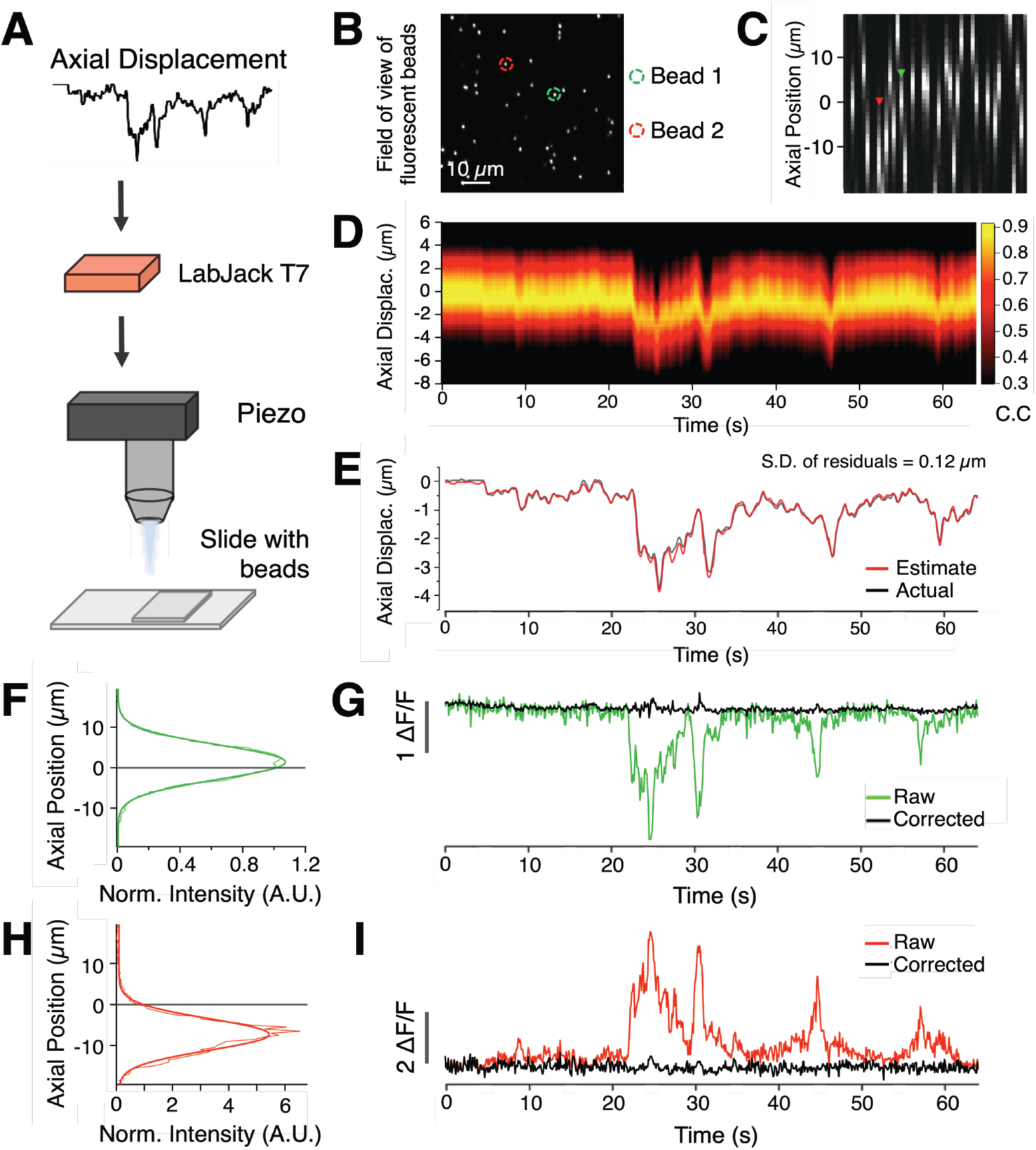
Validating the correction strategy with beads. **A**. Displacement estimation from an *in vivo* experiment is reproduced in the movements of the objective using a piezo controlled focus system while imaging 1 µm beads embedded in agarose to simulate synapses with no activity and no movement of the sample. **B**. Single plane of an example field of beads. Two beads are highlighted as examples used later. **C**. Axial profiles of beads in same field of view as B. **D**. Correlation map between the slices of the reference stack (y-axis) and the time-series (x axis). **E**. Estimate of z-displacement extracted from the correlation results (red trace) closely follows the imposed movement. Standard deviation of residuals was 0.12 µm. **F**. The axial profile of an example bead (green ROI in B), with its peak above the imaging plane (grey line). The continuous line describing the measurements is a Gaussian with sd = 5.2 μm. Thin coloured line shows raw intensity values at each slice, thick coloured line is a fitted Gaussian. Signal was normalised to the intensity at 0 µm displacement **G**. Intensity trace of the same bead before (green) and after correction (black), eliminating shifts in intensity caused by axial displacement. **H** and **I**, as for F and G, for another example bead with its peak below the imaging plane (red ROI in B). The continuous line describing the measurements in H is a Gaussian with sd = 6.1 μm.

**Figure 4.**
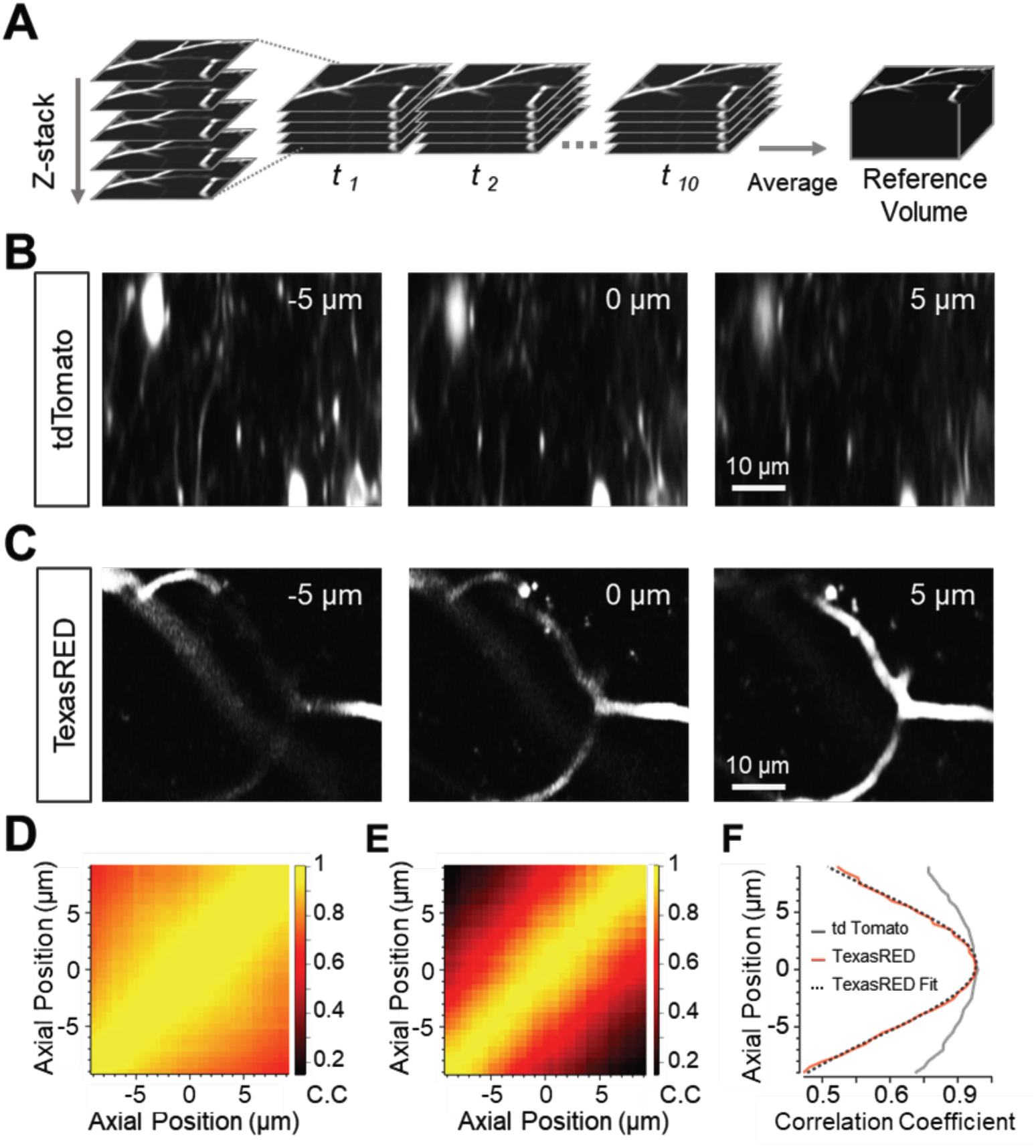
An anatomical marker of z-position: a comparison of tdTomato and dextran-TexasRED in blood vessels. **A**. Schematic showing construction of reference anatomical volume. Multiple stacks are rapidly acquired to enable registration of volumes before collapsing to an average volume. **B**. Two-photon images of VIP interneurons in mouse V1 labelled with tdTomato. Images are example slices from a reference stack acquired as in A at 5 µm above (left panel) or below (right panel) a central plane (central panel). **C**. As in B but with blood vessels labelled with dextran-TexasRED. **D** and **E**. Cross-correlation matrices of an average reference stack with individual slices from a single registered volume for tdTomato (D) and labelled blood vessels (E). **F**. Cross-correlogram of the central slice of the volume with the reference stack for tdTomato (grey) and labelled blood vessels (red). A Gaussian (dotted line) is fitted to the TexasRED correlogram (sd = 8.9 µm) and the maximum taken as the estimated axial location of this slice. Note that the modulation of the correlation coefficient as a function of displacement is much larger for capillaries filled with TexasRED compared to the neurons labelled with tdTomato.

### Workflow for correction of fluorescence signals

To disambiguate synaptic activity from changes in fluorescence caused by z-motion artefacts we constructed a *post hoc* analysis pipeline to automatically correct fluorescence signals from ROIs corresponding to presynaptic structures. The basic elements in this pipeline and their interactions are summarised in Fig. 1. A reference volume containing structural information is required in addition to time-series data acquired in a single plane within that volume. Both the reference volume and time-series comprise two colour channels that separate signals from an anatomical marker and the reporter of neural activity. Here we used a reporter of synaptic activity that fluoresces in green (SyGCaMP6f, Channel 1) and a marker of blood vessels fluorescing in red (Dextran-TexasRED, Channel 2). The steps in the correction procedure applied to these image sets are described below while the rationale is explained and validated in Results.

All image processing and analysis were performed using custom software written in Igor Pro (Wavemetrics, Portland, OR, U.S.A.) including the analysis package SARFIA (Dorostkar *et al*., 2010). These procedures are available on Github (https://github.com/lagnadoLab/zCorrect) and will also operate on activity data matrices produced by other packages commonly used to analyse multiphoton experiments, such as Suite2P (Pachitariu *et al*., 2016) or NMF pipelines (Maruyama *et al*., 2014; Pnevmatikakis *et al*., 2016). The description that follows is designed to provide the information required to code similar pipelines in other languages.

#### i) Collection of a reference volume

Before imaging the activity reporter, a reference volume was acquired that spans the plane which will be imaged during the actual experiment and extending beyond any z-displacement likely to occur during the experiment. We typically acquired a volume spanning 10 µm in either direction from the intended imaging plane, in 0.5 µm steps. While smaller z-steps can result in a more accurate estimation of location, the acquisition time increases. Conversely, larger intervals can be acquired more quickly, but lead to poorer estimations. We usually acquired 10 volumes in quick succession that were then registered in 3D and averaged to produce a high-quality reference volume for subsequent estimates of z-motion. Acquisition of the standard reference volume took 40 s when imaging at 0.1 s per frame.

#### ii) Image pre-processing and estimation of axial displacement

The time-series data and reference volume were automatically split into their two imaging channels. In the time-series, Channel 2 is smoothed in x and y using a Gaussian filter (s.d. = 3 pixels, ∼0.6µm) to reduce fluctuations in intensity of the vessels as red blood cells pass through. Using 2D cross-correlation, we then found the position within the reference stack that most closely matched the average of the time-series. This was defined as the position of zero displacement. This step corrected for any drift in x and y that occurred between acquisition of the reference stack and the time-series and ensured that ROIs defined in the time-series corresponded to the same pixels in the reference stack. Realignment introduced artefacts at the borders of images that influenced the estimation procedures so the largest shifts in the x and y dimensions during each realignment step were used to crop the borders of both the time-series and reference volume. Each frame in Channel 2 of the time-series was then cross-correlated with all the frames in the reference volume to estimate axial motion as a function of time during the recording, one of the three relationships required to correct for intensity changes of the reporter during z-motion (relation 3 in Fig. 1). Estimates of z-displacement during normal behaviour of the mouse were accurate to within ∼0.1 µm (Fig. 3).

#### iii) Identification of ROIs, extraction of time-series and axial intensity profiles

ROIs were identified using the SARFIA analysis suite (Dorostkar *et al*., 2010). Briefly, a standard deviation projection of Channel 1 of the realigned time-series was made and then transformed through a Laplace operator to identify enclosed structures that fluctuated in intensity. These structures had to pass a minimum area threshold to be counted as an ROI, usually of 0.89 µm^2^. Pixel intensity values for each ROI were then averaged on a frame-by-frame basis to generate the “raw” (uncorrected) signal for each ROI in the time-series (relation 1 in Fig. 1). The same ROI masks were then applied to Channel 1 of the reference volume to obtain the relation between ROI intensity and z-displacement (relation 2 in Fig. 1).

An important advantage of using SyGCaMP rather than a cytoplasmic reporter to detect synaptic activity is that the background signal is lower because the neuropil is not fluorescent, resulting in an improved signal-to-noise ratio (Dreosti *et al*., 2009). Any scattered signal from nearby synapses was calculated on an ROI-by-ROI basis by averaging the signal in a “halo” of pixels extending ∼1.5 times the width of the ROI, excluding any that fell within another ROI. The relative intensity of this background signal was then reduced by a ‘contamination ratio’ equivalent to the space occupied by the ROI itself, estimated at 0.5 here and elsewhere (Kerlin *et al*., 2010; Peron *et al*., 2015), after which it was subtracted from the raw signal. The same procedure was used to subtract the background from ROIs defining synapses in Channel 1 of the reference volume.

Activity traces were expressed as relative changes in fluorescence by dividing the change in fluorescence by the baseline fluorescence (ΔF/F_0_). The baseline F_0_ was usually computed as the mode of the entire activity trace when this was close to the minimum of that trace. Where it was necessary to display negative shifts in intensity, the baseline was defined as the minimum value in the time-series (e.g Fig. 2E, top graph, 3G and 6D bottom graph). This approach to displaying the change in the fluorescence signal F represent the relative change in intensity at the same magnitude regardless of whether the modulation is positive or negative. For the purpose of finding a zero ‘baseline’ in these cases, the mode was taken after normalisation and subtracted.

#### iv) Correcting fluorescence changes caused by z-displacement

The intensity profile of each ROI in z (relation 2 in Fig. 1), measured in the absence of locomotion or visual stimulation, provided the correction factor applied to each frame of the time-series, according to the z-position of that frame. To prevent the introduction of noise into the correction, the measured z-profile was fitted with a continuous function. Although the PSF function of an optical microscope is most often described as a Gaussian (Zhang *et al*., 2007; Small & Stahlheber, 2014), we described the “ROI-spread-function” as a Moffat function of the form:

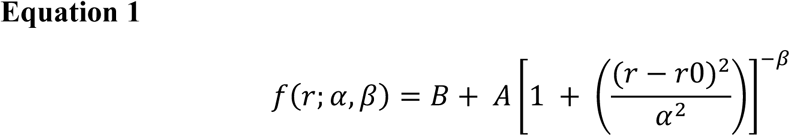

Where ***r*** is the axial location. **A** is the maximum amplitude and **B** is the baseline. We also introduce the parameter ***r0*** to allow the axial location of the synapse centre to vary. To form the probability density distribution, **A** is usually a normalising factor equal to 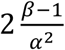. Since we are fitting a function with the same shape as this distribution but with variable amplitudes, we represent this as a free parameter, **A. α** and **β** are so called ‘seeing parameters’ that account for photon scattering through the medium between the object observed and the optics of the detecting device. In astronomy, **α** and **β** reflect aberrations introduced by turbulence in the atmosphere when imaging stars (Moffat, 1969; Racine, 1996; Starck *et al*., 2007). In multiphoton imaging *in vivo* the “seeing parameters” reflect non-uniform scattering through the brain and cranial window (Ji *et al*., 2012). Moffat functions provided significantly better descriptions of the z intensity profile of beads and synapses compared to a Gaussian (Fig. 5). The Igor procedures we provide are easily modified to use a Gaussian or other user defined function, or a simpler smoothing spline (see procedure “correctQA”).

**Figure 5.**
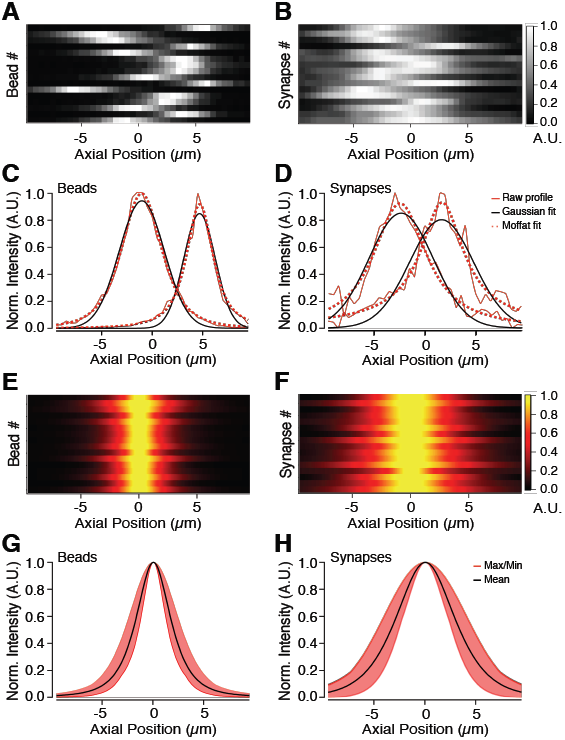
Moffat functions provide a better description of axial intensity profiles than Gaussians. **A** and **B**. Axial profiles for a selection of 16 fluorescent beads (A) or synapses (B) from a single field of view. Each row represents an ROI and has been normalised to the maximum for display purposes. **C**. Raw axial profiles of 2 example beads (red lines), together with best-fit Gaussian (black) and Moffat (red dotted) functions. For the left-hand profile, Gaussian s.d.=3.15 µm and Moffat function has α =3.58 and β=1.97. For right-hand profile, Gaussian s.d.=2.2 µm and Moffat function α=1.9 and β=1.35. Note that the peak and tails of the profile is underestimated by the Gaussian fit, but not the Moffat function. **D**. As in C but for 2 example synapses. For the left-hand profile, Gaussian s.d.=2.5 µm and Moffat function has α =3.3 and β=1.09. For right-hand profile, Gaussian s.d.=3.7 µm and Moffat function α=1.7 and β=0.49. Again, a Moffat function provides a better description than a Gaussian. **E** and **F**. Fitted axial profiles of the same beads (E) and synapses (F) as in A and B with peaks realigned to the centre, showing the homogeneity of beads compared to synapses. **G**. Average Moffat function fitted to profile of beads has α=2.73 and β=1.59 (black line). The red shaded area shows the variance. **H**. Average Moffat function fitted to profile of synapses has α=5.95 and β=2.13 (black line).

The fitted Moffat function is then normalised by the fluorescence intensity of that ROI at the starting position of the time-series, as shown in the examples in Fig. 3F and H. The function relating z-displacement to a relative change in signal for each ROI (relation 2 in Fig. 1) is then used as a look-up table such that the z-position of each frame (relation 3 in Fig. 1) can be converted into a vector of correction factors for each ROI (relation 4 in Fig. 1). A division factor of 1 is found if no displacement occurred within a time frame. Shifts away from the plane in which the ROI is in best focus give values of less than 1, while shifts towards it give values of more than 1. The background-corrected signal within each ROI (relation 1 in Fig. 1) is then divided by the correction vector (relation 4 in Fig. 1) to produce the final signal in which z-motions have been compensated (relation 5 in Fig. 1).

#### v) Curation of activity traces

Following correction, a number of criteria were imposed to remove ROIs that could not be confidently considered synapses. *i)*. ROIs with z-profiles containing more than one peak indicated a contaminating signal from a second synapse and were rejected. *ii)*. Only synapses that were described well by the Moffat function were passed, as assessed by a threshold chi-square value of 0.6, with chi-square normalised to the maximum of each fit. This criterion was set following inspection of the distribution of chi-square values for four fields of view. *iii*). Profiles with a full width half maximum (FWHM) of <4 µm or >10 µm were also excluded as these ROIs are unlikely to be single synapses with these spatial dimensions. *iv*). If the axial displacement caused the expected ROI signal to fall by more than 90% that ROI was also excluded, to avoid corrections obtained by dividing by a value close to zero (step 5 in Fig. 1). These selection criteria can be modified by the user within the software we provide.

### Correlating synaptic activity with running speed

Spearman’s rank correlation coefficients were computed between each ROI’s normalised activity trace (ΔF/F_0_) and running speed at each frame after smoothing with a moving average time window of 3 time frames (∼275 ms). In order to determine if the computed correlation was statistically significant, a bootstrap technique was employed in which running speed traces were circularly shifted from a random origin before recomputing correlation with activity as described before (Dipoppa *et al*., 2018). This process was repeated 1000 times, from which a probability distribution of correlation coefficients was obtained, allowing a p-value to be calculated for the coefficient value from the actual trace. The threshold for significant positive or negative correlation was set at p<0.05.

## Results

### An example of the problem: imaging synaptic activity in awake behaving mice

Multiphoton imaging of awake mice has revealed strong modulation of signals in the primary visual cortex (V1) according to the locomotor state of the animal. Many principal cells and VIP (vasoactive intestinal polypeptide) expressing interneurons show increased basal activity and larger responses to specific visual stimuli when the animal is running compared to when it is still (Niell & Stryker, 2010; Ayaz *et al*., 2013; Saleem *et al*., 2013; Erisken *et al*., 2014; Fu *et al*., 2014; Zhang *et al*., 2014; Pakan *et al*., 2016; Dipoppa *et al*., 2018; Khan & Hofer, 2018). A small subset of VIP interneurons are, however, inhibited during locomotion (Dipoppa *et al*., 2018), indicating that changes in behavioural state have differential effects on different sub-circuits within V1. A key component of these circuits are the synapses that transfer signals between neurons, but these cannot be assumed to be a simple reflection of somatic activity. For instance, the effects of different neurons may be weighted differently simply because they provide different numbers of synaptic outputs. Further, neuromodulators generating changes in internal state, such as acetylcholine and noradrenaline, often act specifically on calcium channels in the presynaptic bouton. It is therefore important to measure synaptic activity directly to understand how changes in behavioural state alter sensory processing.

Using mice free to run on a polystyrene treadmill, we imaged SyGCaMP6f expressed in VIP interneurons in V1 (VIP; Fig. 2A and B), while animals were presented with either uniform grey illumination or a full field drifting grating stimulus (spatial frequency 0.04 cycles per degree; temporal frequency 2 Hz; 100% contrast). SyGCaMP6f is a fusion of the calcium reporter GCaMP6f and the synaptic vesicle protein synaptophysin and is localized to synaptic boutons where it reports the calcium transients that activate neurotransmitter release (Dreosti *et al*., 2009). Two such boutons at different depths within layer 2/3 are highlighted by red and green circles in Fig. 2B. Being small relative to the PSF of the microscope, the brightness of these structures changed significantly when the focal plane was altered by 5 µm, as shown by Fig. 2C and D. As imaging depth increased, the intensity of ROI 1 (red) decreased, while ROI 2 (green) increased (Fig. 2D).

The change in the signal from these ROIs over a 60 s period is shown in Fig. 2E and compared to locomotor activity measured as running speed. It can be seen that two strong bouts of running (shaded) were associated with a decreased signal in ROI1 and increased signal in ROI 2. A visual stimulus consisting of a drifting grating covering the whole visual field was also applied during periods shown by blue bars; the apparent change in synaptic calcium signals were surprising because such a wide-field stimulus normally has little effect on the activity of VIP interneurons measured either in the absence or presence of locomotion (Pakan *et al*., 2016; Dipoppa *et al*., 2018). Doubts over whether the observed SyGCaMP6f signals truly represent changes in synaptic activity were compounded by the observation that the signals in the two ROIs changed in the same direction in response to a downward displacement of the focal plane in the absence of locomotion (Fig. 2D). The concern then, is that when the animal runs there is also a shift in focal plane and therefore a modulation of fluorescence that is not caused by neural activity.

### Strategy for correcting axial motion artefacts

To disambiguate z-motion artefacts from changes in synaptic activity we used images acquired through the multiphoton microscope to estimate three relationships, as summarised in Fig. 1. i) The fluorescence signal in each ROI as a function of time, while imaging a single plane. ii) The signal in each ROI as a function of z-displacement – the axial profiles. iii) The z-position of each frame in the time-series. With knowledge of these relations, the signal in each ROI was corrected to obtain an estimate of activity *independent* of z-position.

To calculate z-position as a function of time (relation 3 in Fig. 1), we first created a 3D reference volume in two colour channels: the green channel collected the signal from SyGCaMP6f and the red channel collected an inactive anatomical marker – a red fluorescent dextran filling blood vessels (see Methods). The same red and green channels were then recorded during the acquisition of the experimental time-series. An estimate of the z-position for each time frame in the time-series (relation 3 in Fig. 1) was obtained by cross-correlating each frame with a reference volume, using the anatomical marker in the red channel. These reference volumes typically spanned a depth of 10-20 µm, sampled in steps of 0.5 µm and acquired just before the experimental time-series. The green reference volume allowed us to estimate how the intensity of each ROI varied as a function of z (relation 2 in Fig. 1). Relations 2 and 3 then allowed us to calculate a “correction vector” for each ROI, describing the relative change in signal caused by z-displacements during the recording, independent of activity (relation 4 in Fig. 1). Each raw signal trace (relation 1 in Fig. 1) was then divided by the corresponding “correction vector” to correct for z-motion artefacts (relation 5 in Fig. 1). These various steps are validated below.

### Testing correction strategy using beads

To assess how accurately the z-position of structures the size of synapses might be estimated by cross-correlation with a reference stack (Fig. 1), we carried out simulation experiments using green fluorescent beads. Beads were 1 µm in diameter, embedded in agarose and covered with three coverslips of #1 thickness, as was used for the cranial window on a mouse (Fig. 3A and B; see Methods). These simulations removed the real-life complications of unpredicted movement and activity-dependent changes in fluorescence. Instead, defined changes in the focal plane relative to the beads were imposed by mounting the objective on a piezo motor that was driven by a voltage signal varying in time to simulate z-displacement during an *in vivo* experiment. An example field of view is shown in Fig. 3B and the axial profiles of each ROI through a reference volume in Fig. 3C. The spectrogram in Fig. 3D shows the correlation coefficient calculated when each frame in the time-series is cross-correlated with each frame in the reference volume. Where there is no displacement from the central imaging plane, the correlation coefficient value is at a maximum with the central plane in the reference volume.

The relation between z position and correlation coefficient (calculated against the reference volume) could be described with a Gaussian function (Figs. 3F and H), from which interpolated estimates of the z shift could be made with a resolution significantly greater than the interval between z slices sampling that volume (usually 0.5 µm). The actual z-displacement of the beads relative to the focal plane over the course of this timeseries is plotted as the grey trace in Fig. 3E, where it is compared with the estimate of z-location over time (red trace). The agreement is very good over the range of displacements in this example (up to 4 µm): the standard deviation of the residuals was 0.12 µm. Similar accuracy in estimating z-displacements of fluorescent beads was observed for deviations of up to 8 µm, using a reference volume covering 10 µm above and below the imaging plane.

The intensity profile in z for the two beads in Fig. 3F and H represents a convolution of the actual z-profile of the beads and the axial PSF of the microscope. A Gaussian was therefore fitted to each profile. From the Gaussian fits to these profiles, we calculated the *expected* relative change in the fluorescence signal within each ROI as a function of imaging depth and then divided by this value to calculate the signal expected in the absence of z-motion. For instance, if a shift of +3 µm was estimated for a given time frame and the model informs us that would lead to a 10% decrease in intensity compared to baseline, the acquired value is 0.9 times that at the original imaging plane. We can then reconstruct the intensity at the intended imaging plane by dividing the ROI’s intensity by a correction factor of 0.9. Fig. 3G and I show the raw fluorescence signal in these two ROIs before (coloured traces) and after correction (black traces). The procedure completely removed fluctuations in fluorescence caused by z-displacements.

In some cases, the magnitude of the shifts was sufficient to move the imaging plane to a position where the ROIs’ intensity had fallen close to background so that information about the intensity of that ROI was missing or unreliable. The ability to reconstruct the true fluorescence therefore depended on the depth of the reference volume relative to z-displacements, the location of the structure in z, and the axial PSF of the microscope. Structures become “lost” if their centre is towards the edge of the reference volume and a z-displacement moves them further away from the focal plane. Excluding these cases, ∼95% of fluorescent beads in three separate fields of view were corrected back to a static baseline (15/16;12/13;18/18) using different axial displacement inputs ranging up to 8 µm.

### Blood vessels as an anatomical marker for *in vivo* imaging

To apply the correction strategy described in Figs. 1 and 3 to images acquired *in vivo* we required an anatomical marker of z-position that was unaffected by neural activity and which could be imaged simultaneously with the green emission from SyGCaMP6f. We first tested a fluorescent protein that emits in the red part of the spectrum by co-expressing tdTomato in the cytoplasm of VIP interneurons, thereby labelling the somas and dendrites (Fig. 4B). Although tdTomato is often used to mark neural structures (Kerlin *et al*., 2010; Fu *et al*., 2014; Besser *et al*., 2015; Dipoppa *et al*., 2018), the correlation coefficients between images and a reference stack were not strongly modulated by displacements in z (Fig. 4B and D). The relatively high degree of similarity between images at different depths reflects the fact that a cytoplasmic marker labels many fine structures throughout the volume (Fig. 4B), leading to low SNR because of scattered light from out of focus neuropil. Normalised correlation values varied between 0.81 and 0.96 for displacements of ± 5 µm (Fig. 4F).

An ideal anatomical marker would label compartments with a spatial structure that varies sufficiently in z to distinguish frames at small distances and provide images of high contrast and low noise. We tested whether blood vessels might provide such images by filling them with dextran-TexasRED introduced into the circulation. This water-soluble dye fills blood vessels but cannot cross the blood brain barrier, producing images of high spatial contrast. TexasRED can be readily excited under two-photon using wavelengths that are also effective at exciting GFP (920 nm) and consequently syGCaMP6f (Fig. 4A and C). It is also simpler to use than virus injection, only requiring a sub-cutaneous injection 15 mins prior to an experiment. Applying the same cross-correlation procedure to images of blood vessels produced much deeper modulation of correlation values as a function of z-displacement compared to tdTomato (0.3 to 0.97; Fig. 4 E and F) and, therefore, more accurate estimates of z-displacement.

### Modelling the z-profile of synapses

With knowledge of the z-position of the imaging plane through a time-series, it might in principle be possible to correct for z-motion artefacts by deconvolution using the PSF of the microscope measured in 3D. In ideal conditions, the PSF is uniform across the field of view and can be defined by a mathematical model of diffraction, when it will have axial and radial symmetry. In practise, the measured PSF will not be symmetrical because of aberrations in the optical system (Ji *et al*., 2012). The PSF of the microscope can be measured by 3D imaging of diffraction-limited fluorescent beads, but this will still not be adequate for correction of images collected *in vivo* because of the optical aberrations introduced by the cranial window and the brain tissue itself (Ji *et al*., 2012; Wang & Ji, 2012; Ji, 2017). Crucially, these aberrations vary as a function of x-y-z position in the brain and depend on the properties of the cranial window as well as the exact angle of this window relative to the optical axis of the microscope. One approach to overcoming these problems is to measure aberrations across the field of view (Schwertner *et al*., 2004*b*, 2004*a*) and to then compensate for these using adaptive optic elements, such as spatial light modulators, introduced into the excitation path (Wang *et al*., 2010; Ji *et al*., 2012; Wang & Ji, 2012). The use of adaptive optics is technically demanding and, to our knowledge, has yet to be validated for compensations due to z-motion of the brain. We therefore took a more empirical approach in which we measured the axial intensity profile separately for each ROI defining a synapse, which we termed the “ROI-spread-function” (RSF).

Examples of the RSF for a selection of beads imaged through a window similar to that used for *in vivo* imaging is shown in Fig. 5A and C and can be compared with the RSF of synapses in VIP interneurons expressing SyGCaMP6f synapses shown in Figs. 5B and D. In both cases, the best-fit Gaussians describing the whole RSF failed to account for the gradual tailing-off of intensity away from the focal plane while also underestimating the peak amplitude (black curves in Fig. 5C and D). A significantly better description of the RSF was obtained by using a Moffat function (broken red lines in Fig. 5C and D; Equation 1 in Methods). Such functions have long been used to correct for aberrations in images of astronomical objects acquired through telescopes (Moffat, 1969; Racine, 1996; Starck *et al*., 2007). Fig. 5E and F show the same beads and synapses as in A and B described centred to compare profiles across ROIs and Fig. 5G and H show the averaged RSF described as Moffat functions. The overall variability of profiles between beads was, unsurprisingly, quite small (Fig. 5G), with an average full-width half-maximum (FWHM) of 3.8 µm (black curve). Synapse profiles on average were broader and more varied in size (Fig. 5H), with an average FWHM of 6.6 µm (black curve), reflecting the additional aberrations introduced by imaging through brain tissue. Using a Moffat function to describe the z intensity profile of each synapse provided a simple method of allowing for the non-uniform aberrations across a field of view.

### Correcting for brain movement *in vivo*

We now apply the tools described above to *in vivo* imaging experiments of the type described in Fig. 2, imaging synaptic activity in VIP interneurons of mouse V1 (Fig. 6A). Just before acquisition of the experimental time-series in one plane, we acquired reference volumes imaging SyGCaMP6f (to calculate the RSF of each synapse) and dextran-TexasRED (to subsequently calculate z displacements during the time-series), as shown in Fig. 6B. Axial motion was estimated by cross-correlation and, as anticipated, locomotor behaviour was associated with shifts in imaging depth (Fig. 6C). Displacements of up to 5 µm were commonly observed, but never any larger than 8 µm. Shifts were typically in the negative direction, meaning the brain moves towards the objective and the imaging plane becomes deeper when the animal is running.

**Figure 6.**
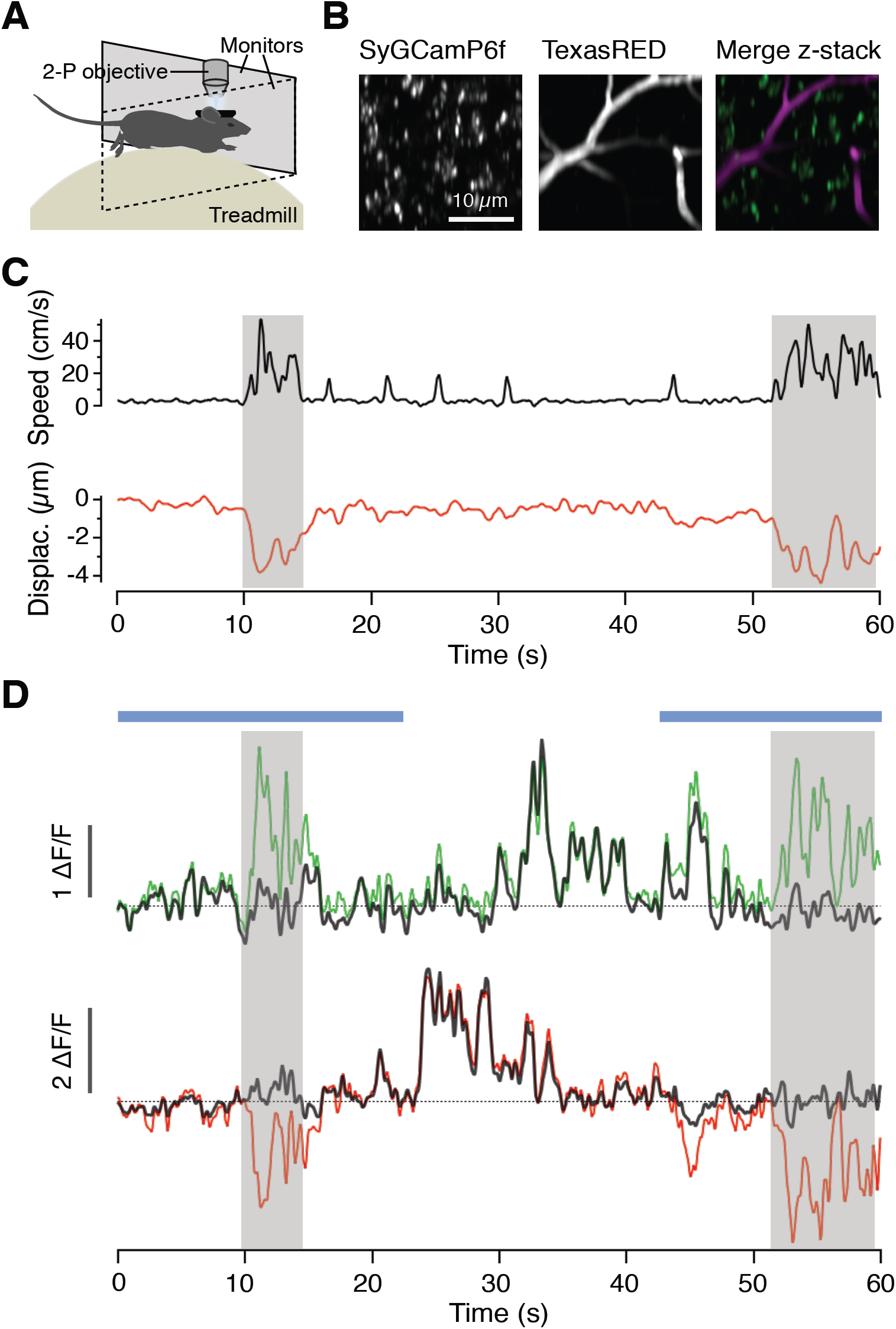
Correcting z-motion artefacts when measuring synaptic activity *in vivo*. **A**. Two-photon (2-P) recording in awake behaving mice, as in Figure 2A. **B**. Two-photon images (average of 50 frames) with synapses labelled with SyGCaMP6f recorded in parallel with blood vessels labelled with dextran-TexasRED. **C**. Running speed (top, grey trace) is associated with axial displacement of the focal plane (bottom, red trace). **D**. Green and red traces are uncorrected activity traces acquired from two example synapses. The corrected traces are shown in grey and these no longer exhibit changes in fluorescence associated with locomotor bouts. Blue bars show two periods of application of a full-field grating stimulus (0.04 cycles per degree, 2 Hz, 100% contrast).

The red and green traces in Fig. 6D show the uncorrected signals from the two synapses featured in Fig. 2 and the dark grey traces show the effects of correcting for z motion by using the RSF as a look-up table to find the appropriate correction factor for each time point. Before correction, the activity of the red synapse appeared to be positively correlated with locomotion (correlation coefficient of 0.32; p = 0.0007) while the red synapse appeared to be inhibited (correlation coefficient of −0.46; p = 0.002). Correcting for changes in fluorescence predicted by the measured z motions and RSF removed modulations of signals during locomotor bouts (black traces) indicating that both were artefacts wholly reflecting z motion without any contribution from neural activity. At other times with little or no axial displacement, the signals from the two synapses were uncorrelated and they were not altered significantly by the correction. The correction procedure compensated for changes in the signal that were correlated between synapses and locomotion but did not alter synaptic signals that were not correlated in the absence of z motion. This observation provides strong evidence that fluorescence signal reflecting changes in synaptic activity were disambiguated from changes in the signal caused by movement of the focal plane.

An overview of the impact of correcting for z motion on signals measured across a population of synapses from VIP interneurons is provided in Fig. 7. When measured at the soma using a cytoplasmic calcium reporter, the activity of the vast majority of VIP neurons is strongly correlated with locomotion and in >90% this correlation is positive (Fu *et al*., 2014; Pakan *et al*., 2016; Dipoppa *et al*., 2018). A different picture emerged when we investigated the apparent correlation between synaptic activity and running speed and the effects of z-motion artefacts. The distribution of correlation coefficients in 1312 VIP interneuron synapses in 22 fields of view in 2 mice are shown before correction in Fig. 7A and after correction in Fig. 7B (all imaged in layer 2/3). Before correction only 38% of VIP synapses appeared to be significantly modulated during locomotion and after correction 32%. Before correction, the majority of synapses that appeared to be significantly affected by locomotion were inhibited (298/497 = 58%), while after correction there was a shift in the distribution towards more positive correlations (279/422 = 63%). These results indicate that somatic activity in VIP interneurons cannot be assumed to be mirrored in a simple way in the output from the same population of neurons, underlining the importance of monitoring synaptic activity in addition to somatic calcium signals.

**Figure 7.**
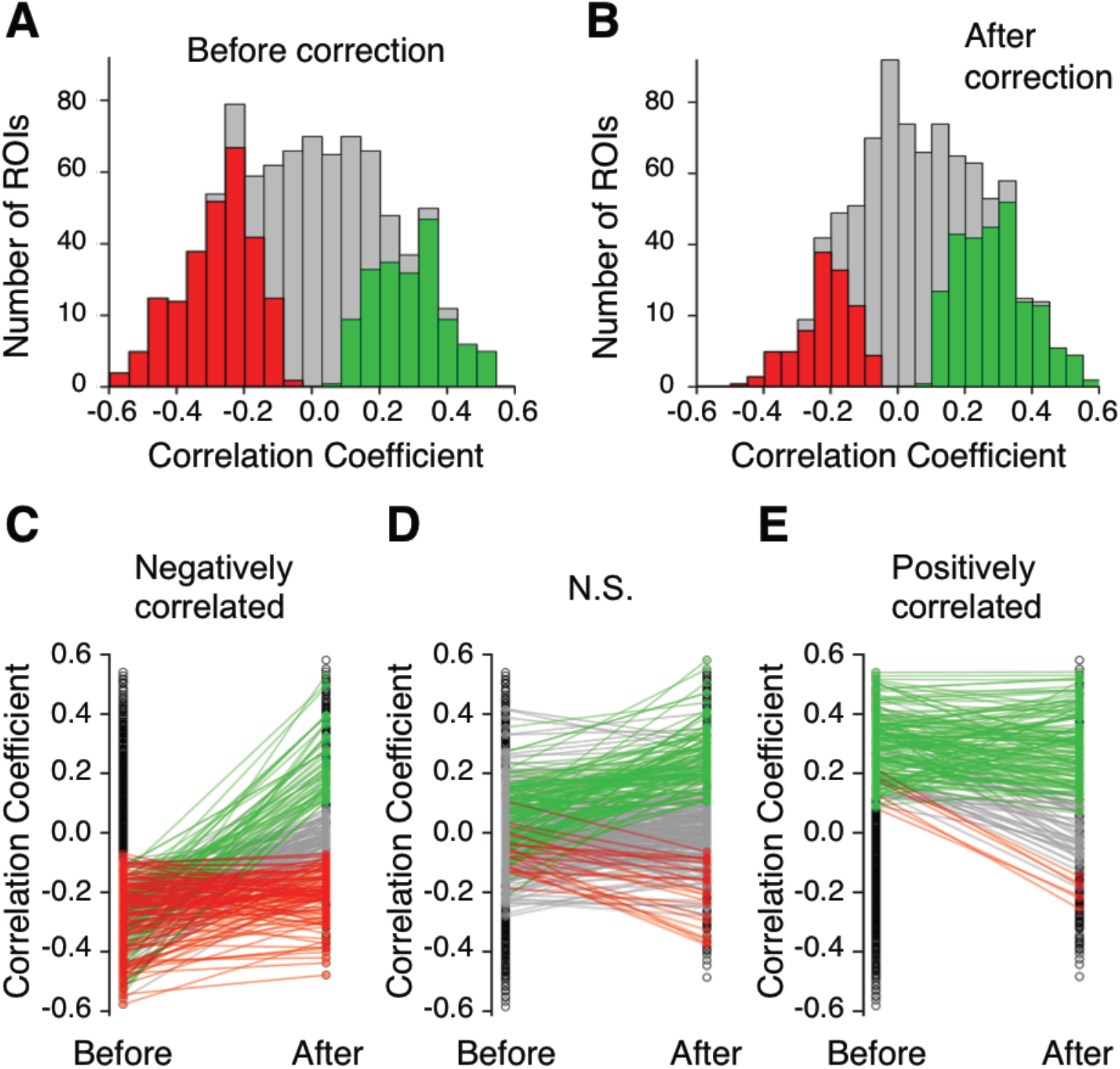
The impact of correcting for z motion on signals measured across a population of synapses. **A**. Distribution of correlation coefficients between synaptic activity trace and running speed, before correction for z-motion. Results collected from 1321 synapses in two mice expressing SyGCaMP6f in VIP interneurons in V1 (layer 2/3). The majority of synapses show some correlation with locomotion. Coloured bars represent synapses that appear significantly correlated, either positively (green) or negatively (red). **B**. Distribution of the same synapses after correction. The majority of synapses now show no significant correlation and a greater proportion are positively correlated. **C-E**. The destination of synapses with significant negative (C), not significant (D), or significantly positive (E) correlation coefficients to running speed prior to correction. Coloured lines and markers show the pairwise correlation coefficient changes for each ROI, showing a shift to positive (green), negative (red) or not significant (grey) correlations with running speed following correction.

To understand further the impact of correcting z motion artefacts, we split our sample of VIP-positive synapses into three groups: those that were positively or negatively correlated with locomotion and those that were not significantly modulated. The effect of the correction procedure on the correlation identity of these three groups is shown in Fig. 7 C-E. Black circles represent the overall distribution of correlation coefficients, while coloured points identify them as not significant (NS, grey), or significantly negative (red) or positive (green): the lines connect measurements for the same synapse before and after correction. Of the 289 synapses that appeared to be negatively correlated before correcting for z-motion artefacts, only 40% retained this identity after correction. In fact, 18% were revealed to be more strongly *activated* during locomotion. Synapses that appeared to be positively correlated before correction were far more likely to have been identified correctly (138/208, ∼66%; Fig. 7E, green lines) with only 3% being revealed to be negatively correlated (6/208, 3%; red lines). The vast majority of synapses that showed no significant dependence on locomotion before correction also showed none after correction (263/374, ∼70%; Fig. 7D, grey lines), while ∼24% were subsequently identified as positively correlated (89/374, green lines) and only ∼6% as negative (22/374, red lines). The overall picture is one in which z motion artefacts can appear to modulate synaptic activity during locomotion: correcting these artefacts altered the apparent dependence of activity on locomotion in 27% of the 1321 synapses.

## Discussion

Here we have described a new and relatively simple approach to identify and remove axial motion artefacts when imaging activity in small neural compartments such as synapses. The method accurately measures displacements of the imaging plane over a time-series and we demonstrate that it can be used to correct activity traces acquired from populations of synapses imaged in the visual cortex of awake behaving mice (Fig. 6). Conveniently, the method can be used when imaging activity in a single plane on any standard commercial multiphoton microscope without the need for further specialised equipment and should prove useful when imaging synaptic activity using reporters such as SyGCaMPs (Dreosti *et al*., 2009; Dreosti & Lagnado, 2011; Nikolaev *et al*., 2013) or iGluSnFR (Marvin *et al*., 2013, 2018).

One approach to solving the problem of shifts in focus during *in vivo* imaging is to continuously scan a volume, thereby allowing volumetric registration similar to the x-y registration that is already standard when imaging in one plane (Holekamp *et al*., 2008; Deneux *et al*., 2016; Kong *et al*., 2016). However, volumetric imaging can only be performed at a fraction of the temporal resolution of single plane imaging, and this represents a serious problem in capturing activity on time-scales that are physiologically relevant. While specialised equipment such as acousto-optical deflectors (Duemani Reddy & Saggau, 2005; Deneux *et al*., 2016) or light-sheet microscopy (Holekamp *et al*., 2008), can facilitate rapid volumetric acquisition it is desirable to find a solution that can be applied to imaging in a single optical plane using a standard multiphoton microscope. One technique is to measure directly the modulation of intensity caused by z-displacements using an anatomical marker that is co-expressed with the reporter of activity, without directly estimating the axial displacement during the time series. Such an approach has been used to, for instance, correct somatic activity using tdTomato (Marinković *et al*., 2019). A disadvantage of using a second fluorescent protein is that it usually requires viral delivery of additional constructs or appropriate transgenic animals as well as potential complications arising from photobleaching. We avoid these issues with a simple sub-cutaneous injection of a fluorescent dextran-conjugated dye that fills small capillaries and found that this provided images of high contrast and estimates of z-displacement superior to those obtained using a second fluorescent protein expressed in the neurons of interest (Fig. 4). Similar fluorescent dyes exist with different spectral properties so this approach can in principle be used during experiments where activity is recorded using reporters emitting at different wavelengths.

Correcting for changes in focal plane was found to alter the apparent correlation between the activity of VIP-positive synapses and locomotor activity in awake mice (Figs. 6 and 7). Somatic activity in VIP interneurons appears to be almost exclusively positively correlated with locomotion (Fu *et al*., 2014; Pakan *et al*., 2016; Dipoppa *et al*., 2018), but we found ∼37% of synapses to be negatively correlated, even after correcting for z-motion artefacts (Fig. 6 and 7). This apparent discrepancy might be explained if few VIP interneurons whose activity is anti-correlated with locomotion deliver their output through larger numbers of synapses than those that are activated during locomotion. Alternatively, this transition could be caused by presynaptic inhibition of presynaptic terminals. Whatever the mechanism, these observations underline the importance of making independent assessments of synaptic activity: the notion that these simply reflect somatic activity is likely too simplistic. Understanding the activity dynamics of the cortex at the level of synapses will be an essential aspect of understanding the dynamics of signal flow through this circuit and its modulation under different behavioural states.

## Funding

This work was supported by a Wellcome Trust Investigator Award (102905/Z/13/Z).

## Notes

https://github.com/lagnadoLab/zCorrect

